# DiscoSnp-RAD: *de novo* detection of small variants for population genomics

**DOI:** 10.1101/216747

**Authors:** Jèrèmy Gauthier, Charlotte Mouden, Tomasz Suchan, Nadir Alvarez, Nils Arrigo, Chloé Riou, Claire Lemaitre, Pierre Peterlongo

## Abstract

We present an original method to *de novo* call variants for Restriction site associated DNA Sequencing (RAD-Seq). RAD-Seq is a technique characterized by the sequencing of specific loci along the genome, that is widely employed in the field of evolutionary biology since it allows to exploit variants (mainly SNPs) information from entire populations at a reduced cost. Common RAD dedicated tools, as *STACKS* or *IPyRAD*, are based on all-versus-all read comparisons, which require consequent time and computing resources. Based on the variant caller *DiscoSnp*, initially designed for shotgun sequencing, *DiscoSnp-RAD* avoids this pitfall as variants are detected by exploring the De Bruijn Graph built from all the read datasets. We tested the implementation on RAD data from 259 specimens of *Chiastocheta* flies, morphologically assigned to 7 species. All individuals were successfully assigned to their species using both STRUCTURE and Maximum Likelihood phylogenetic reconstruction. Moreover, identified variants succeeded to reveal a within species structuration and the existence of two populations linked to their geographic distributions. Furthermore, our results show that *DiscoSnp-RAD* is at least one order of magnitude faster than state-of-the-art tools. The overall results show that *DiscoSnp-RAD* is suitable to identify variants from RAD data, and stands out from other tools due to his completely different principle, making it significantly faster, in particular on large datasets.

**License:** GNU Affero general public license

**Availability:** https://github.com/GATB/DiscoSnp

## 1 Introduction

Next-generation sequencing and the ability to generate thousands of genomic sequences has opened new horizons in population genomic research. This has been made possible by the development of cost efficient approaches to obtain sufficient homologous genomic regions, by reproducible genome complexity reduction and multiplexing several samples within a single sequencing run [1]. Among such methods, the most widely used over the last decade is “*Restriction-site associated DNA sequencing*” (RAD-Seq). It uses restriction enzymes to digest DNA at specific genomic sites and sequence the adjacent regions. This approach encompasses various methods with different intermediate steps to optimize the genome sampling, e.g. ddRAD [15], GBS [4], 2b-RAD [22], 3RAD/RADcap [9]. These methods share basic steps: DNA digestion by one or more restriction enzymes, ligation of sequencing adapters and sample-specific barcodes, followed by optional fragmentation and fragment size selection, multiplexing samples bearing specific molecular tags, i.e. indices and barcodes, and finally sequencing. The sequencing output is thus composed of hundreds of thousands of reads originating from all the targeted homologous loci. The bioinformatic steps consist in sample demultiplexing, rebuilding the loci and identifying informative homologous variations. If a reference genome exists, the most widely used strategy is to align the reads to this reference genome and to perform a classical variant calling: Single Nucleotide Polymorphism (SNP) and Insertion-Deletion (INDEL). However, RAD-Seq approaches are mainly used on non-model organisms for which a reference genome does not exist. The fact that all reads sequenced from the same locus start and finish exactly at the same position makes it easier to compare directly reads sequenced from a same locus. To *de novo* build homologous genomic loci and extract informative variations different methods have been developed, *STACKS* [2] and *PyRAD* [3], as well as its derived rewritten version *IPyRAD* (https://github.com/dereneaton/ipyrad), being the most commonly used in the population genomics community.

The main idea behind these approaches is to group reads by sequence similarity in clusters representing each a distinct genomic locus. Sequence variations can then be easily identified and a consensus sequence is built for each locus, since reads start and end at the same position. The key challenge is therefore the clustering part. To do so the classical approach relies on all-versus-all alignments. To reduce the number of alignments to compute, the clustering is first performed within each sample independently, then sample consensuses are compared between samples. Nevertheless the number of alignments to perform remains very large and increases quadratically with the number of reads. Importantly, analysis of RAD data is highly dependent on the chosen method, the sequencing quality and the dataset composition, such as the presence of inter and/or intra-specific specimens or the number of individuals. Thus existing tools allow customization of numerous parameters to fine-tune the analysis. Particularly, both methods have parameters controlling the granularity of clustering: the number of mismatches allowed between sequences of a same locus within and among samples for *STACKS* and the percentage of similarity for *PyRAD*. These have a significant impact on downstream analyses.

We present here an utterly different approach to predict *de novo* small variants from large RAD-Seq datasets taking advantage of the *DiscoSnp++* approach [21, 14]. Initially, *DiscoSnp++* was designed for *de novo* prediction of SNPs and small INDELS, from shotgun sequencing reads, typically whole genome re-sequencing data, without the need of a reference genome. The basic idea of the method is a careful analysis of the *de Bruijn graph* built from all the input read sets, to identify topological motifs, often called *bubbles*, generated by polymorphisms. In this work, we propose an adaptation of the *DiscoSnp++* approach to the RAD-Seq data specificities. After a small proof-of-concept test on simulated data from *Drosophila melanogaster*, we present an application of the *DiscoSnp-RAD* implementation on double-digest RAD-Seq data (ddRAD) from a genus-wide sampling of parasitic flies belonging to *Chiastocheta* species. Using *DiscoSnp-RAD*, the 259 individuals analyzed could be assigned to their respective species. Moreover, within-species analyses focused on one of these species, identified variants revealing population structure congruent with sample geographic origins. Thus, the information obtained from SNPs and INDELs identified by *DiscoSnp-RAD* can be successfully used for population genomic studies. The main notable difference between *DiscoSnp-RAD* and concurrent algorithms stands in the execution time, as it was more than 15 times faster than *STACKS* run as well as *IPyRAD* run.

## 2 Material and Methods

### 2.1 *DiscoSnp-RAD*: RAD-Seq adaptation of *DiscoSnp++*

Originally, *DiscoSnp++* was designed for finding variants from whole genome sequencing data. In the RAD-Seq context, *DiscoSnp++* modifications affect core algorithm modification, as well as result post-processing.

#### DiscoSnp++ basic algorithm

We first recall the fundamentals of the *DiscoSnp++* algorithm, which is based on the analysis of the *de Bruijn Graph* (dBG) [16] which is a directed graph where the set of vertices corresponds to the set of words of length *k* (*k*-mers) contained in the reads, and there is an edge between two *k*-mers if they perfectly overlap on *k* − 1 nucleotides. Small variants, such as SNPs and INDELs, generate in the dBG recognizable patterns called “*bubbles*”. A bubble (Fig.1(a)) is defined by one *start* branching node that has, two distinct successor nodes. From these two children nodes, two paths exist and merge in a *stop* branching node, which has two predecessors.

**Fig. 1.**
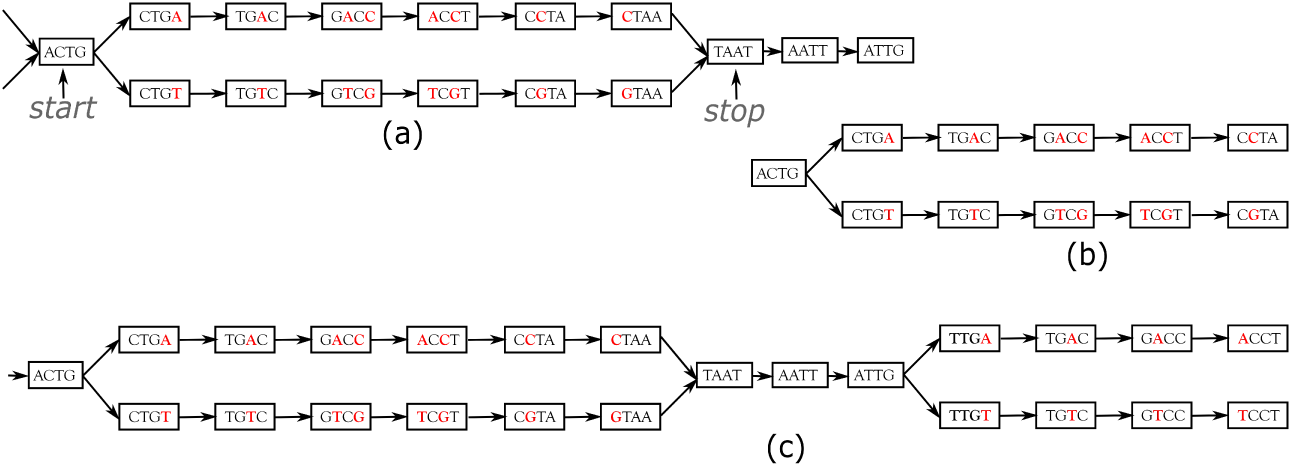
Examples of bubbles generated by SNPs in a toy de Bruijn graph (with *k* = 4). In **(a)** the bubble is complete: this corresponds to a bubble detected by *DiscoSnp++*. Note that *DiscoSnp++* also detects more complicated bubbles generated by short INDELs and/or containing branching nodes themselves (not shown here). In **(b)**, the bubble is truncated: it is composed of a branching node (“*ACT G*”) whose two successors lead to two distinct paths that both have the same length and such that their last two nodes have no successor. Graph **(c)** shows an example of two bubbles from the same locus.

*DiscoSnp++* builds a dBG from all the input read datasets combined, and then detects such bubbles. Sequencing errors or approximate repeats also generate such bubbles, that can be avoided by using of a minimal read coverage threshold to keep a bubble (`-c` parameter), and limiting the type of authorized branching nodes on the two paths (`-b` parameter).

#### A novel RAD-specific bubble model

In *DiscoSnp++*, variants distant from less than *k* bp from a genomic extremity could not be detected, as associated bubbles do not open and/or close. This effect is negligible in the whole genome sequencing context, however, in the RAD-Seq context, sequenced genomic regions are limited to one or a few hundreds of nucleotides (the read size), and thus a large amount of variants are likely to be located at the extremities of the loci. For instance, with reads of length 100bp, and *k* = 31 (which is a classical *k* value), on average 62% of the variants are located in the first or last *k* nucleotides of a locus and cannot be detected by *DiscoSnp++*.

In the RAD-Seq context, all reads sequenced from the same locus start and end exactly at the same position. Thus, variants located less than *k* bp from loci extremities generate what we call *Symmetrically Truncated Bubbles* (Fig.1(b)). Such bubbles start with a node which diverges into two distinct paths that do not meet back, such that both of them cannot be extended because of absence of successor and both paths have exactly the same length.

Additionally, we constrain the last 3-mer of both paths to be identical (contrary to what is shown Fig.1). This avoids confusing an INDEL occurring near the extremity with 3 successive substitutions. Although this prevents the detection of variants as close as 3 bp from a locus extremity, this enables to identify correctly the type of detected variant. Note that this issue is also present in any mapping or clustering based approaches. Detected bubbles (truncated or not) are output as whole sequences in fasta format and all variants are reported in vcf format, along with numerous additional information, such as read coverage, quality and ranking information (see [21, 14]).

#### Post-processing and RAD-specific filtering of predicted variants

We now present how variants predicted from bubble enumerations can be exploited in the RAD-Seq context. Notably, after variant calling, several post-processing and variant filtering steps are usually performed to keep only reliable and informative variants for downstream population genomics analyses. Among those, some RAD-Seq filters apply at the locus level. For instance, for population STRUCTURE analysis input variants should not physically linked on the genomes. Another example is the filtering of sites presenting an excess of heterozygosity in a single locus, potentially resulting from artifactual loci built from several paralogous genomic regions. However, in the *DiscoSnp-RAD* approach, bubble detection does not provide links between variants from the same locus. This is why we developed a post-processing method to cluster predicted variants per locus.

#### Grouping variants coming from the same locus

During the bubble detection phase, several independent bubbles can be predicted for the same locus. For instance, Fig.1(c) shows a toy example of a the dBG graph associated to a locus. In this case, *DiscoSnp-RAD* detects two bubbles, that give no sign of connection. However, *DiscoSnp-RAD* is parameterized to output bubbles together with their left and right context in the graph, which corresponds to the paths starting from each extreme node and ending at the first ambiguity (ie. a node with not exactly one successor).In this case, the two bubbles of Fig.1 are output as 2×2 longer sequences (ACTG**A**C**C**TAATTG/ACTG**T**C**G**TAATTG and TAATTG**A**CCT/ TAATTG**T**CCT) that share at least one *k* − 1-mer (here *k* − 1-mers TAA, AAT, ATT and TTG).

If a given locus contains several variants, each bubble of this locus should share one *k* − 1-mer with at least one other bubble of the same locus. We exploit this property to group all bubbles per locus. For doing so, we create a graph in which a node is a bubble (represented by its pair of sequences), and there is an edge between two nodes if the corresponding sequences share at least one *k* − 1-mer. This is done using *SRC linker* [12]. Finally, we partition this graph by connected component. Each connected component contains all bubbles for a given locus and this information is reported in the vcf file. Note that, clustering is performed only on variants with less than 95 % of missing genotypes, since variants with too many missing data can be numerous and are often non-informative and filtered-out later in downstream analyses.

#### Various filtering options

Importantly, the cluster ID is indicated in the final vcf, in the CHR field, enabling to apply custom filters based on cluster information, as well as any variant level classical RAD-Seq filters (such as the minimal read depth to call a genotype or the minimal minor allele frequency to keep a variant). To apply these filters, we provide a catalogue of scripts, specific to RAD-Seq context (https://github.com/GATB/DiscoSnp/tree/master/scripts_RAD).

In particular, paralogous genomic regions represent a major issue in population genomic analyses as DNA sections arising from duplication events can be aggregated in the same locus and thus, might encompass alleles not deriving from coalescent events. We propose two ways to filter out such paralog-induced variants. The first filter takes advantage of the *rank* value computed for each variant. We have designed this scoring scheme and shown in previous work [21, 14] that approximate repeats are likely to generate bubbles in the dBG but with very low rank values (< 0.2) contrary to real SNPs. By default, *DiscoSnp-RAD* discards all variants with such low rank values. The second filter is inspired by classical RAD-Seq pipelines. It uses the clustering results. It assumes that most of the variants of a same locus built from several paralogous sequences will show a similar pattern of excess of heterozygous genotypes. We propose a script to filter out all the variants of a cluster showing more than *X* % of variants each having more than *Y* % of heterozygous genotypes.

### 2.2 Tests on simulated datasets

#### Simulation protocol

We simulated RAD loci from *Drosophila melanogaster* genome (dm6) by selecting 150bp on both sides of 5,000 PstI restriction sites. Each locus was duplicated in two copies and SNPs were randomly introduced at a rate of 0.5 % in the first copy, and a subset of them (30%) was introduced in the second copy, so that loci present both heterozygous and homozygous SNPs. This process was done five times to mimics distinct RAD data from five individuals, with shared SNPs between them. Finally, 37,101 genomic positions were mutated (3.7 SNPs per locus on average). Forward 150bp reads were simulated on right and left loci, with 1% sequencing errors, with 60X coverage per individual. All reads were exactly aligned, as this is the case in RAD data.

#### Evaluation protocol

For estimating the result quality, predicted variants were localized on the *D. melanogaster* genome and output in a vcf file. To do so, we used the standard protocol of *DiscoSnp++* when a reference genome is provided, using BWA-mem [11]. The predicted vcf was compared to the vcf storing simulated variant positions to compute the amount of common variants (true positive or TP), predicted but not simulated variants (false positive or FP) and simulated but not predicted variants (false negative or FN). Recall is then defined as 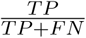 and 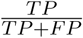.

#### Comparison with other tools

For comparisons, *STACKS* and *IPyRAD* were run on the simulated data. Stacks were generated *de novo* (denovo map.pl), with a minimum of 3 reads to consider a stack (-m 3). Parameters governing the merge of stacks (-m and -M) were fixed to 6, which is coherent with the number of simulated mutations. *IPyRAD* was run using default parameters: a clustering threshold of 0.85 and a minimum depth of 5.

Then, *de novo* tags from *STACKS* and loci from *IPyRAD* were mapped to the *D. melanogaster* genome. For *STACKS*, loci were mapped using GSNAP/GMAP [23]. Genomic coordinates were incorporated in outputs using a script provided in the *STACKS* tools suite, before generating a vcf file with populations module [13]. For *IPyRAD*, loci were mapped to the *D. melanogaster* genome using BWA-mem and variant positions were transposed on the genome positions with a custom script.

### 2.3 Application to real data from *Chiastocheta* species

#### Data origin

Tests on real data were performed on ddRAD reads previously obtained for the phylogenetic study of seed parasitic pollinators from the genus *Chiastocheta* (Diptera: Anthomyiidae). The dataset corresponds to reads from 259 individuals sampled from 51 European populations generated by Lausanne University, Switzerland [19] (data soon available on zenodo.org). Quality scores of reads 2 (average base quality over reads and across samples of 33.98 sd 1.78) were lower than those of reads 1 (average base quality over reads and across samples of 35.15 sd 1.70), which led us to focus the analyses on reads 1. Finally, 300,637,358 reads were used for the study with an average of 1,160,762 reads per individual.

#### Variant prediction and filtering

*DiscoSnp-RAD* was run with parameters `-b 1 -P 5 -D 10`: this authorizes non symmetrically branching bubbles (see [21] for details), searching for at most five variants per bubble, and indels of size of at most 10. Variant predictions (with less than 95% missing data) were clustered and output in vcf format as described above.

Highly heterozygous clusters were filtered out by removing those harbouring more than 50% of heterozygous SNPs in more than 10% of the individuals (`filter paralogs.py`). This latter treatment of predictions is considered as the default process of *DiscoSnp-RAD* outputs. Then, classical filters were applied to follow as much as possible the filters used in the Suchan *et al.* [19]: a minimum genotype coverage of 6, a minimal minor allele frequency of 0.02 and a minimum of 20 samples with a non missing genotype for each variant. These filters remove less informative variants or alleles specific to a small subset of samples. These filters were also applied at intraspecific level in one of the seven sampled *Chiastocheta* species, i.e. *C. lophota*, on the same *DiscoSnp++* output, the only difference being the minimum number of samples to keep a variant set to 2, to remove sample specific variants.

#### Population genomic analyses

The species genetic structure was inferred using STRUCTURE v2.3.4 [17]. This approach requires unlinked markers, thus only one variant by locus, randomly selected, has been kept. The STRUCTURE analysis was carried out using two datasets – the first with both SNPs and INDELs and the second with SNPs only, Simulations were performed with genetic cluster number (*K*) set to 7, corresponding to the seven species described in [19]. We used 20,000 MCMC iterations after a burn-in period of 10,000. The output is the posterior probability of each sample to belong to each of seven possible clusters.

#### Phylogenomic analyses

Maximum likelihood (ML) phylogenetic reconstruction was performed on a whole concatenated SNP dataset using GTRGAMMA model with the acquisition bias correction [10]. We applied rapid Bootstrap analysis with the extended majority-rule consensus tree stopping criterion and search for best-scoring ML tree in one run, followed by ML search, as implemented in RAxML v8.2.11 [18].

## 3 Results

### 3.1 Results on simulated data

*DiscoSnp-RAD* was first run on a simple simulated RAD-Seq dataset in order to provide a proof of concept of the approach and to compare it with the other clustering approaches. This simple experiment shows that *DiscoSnp-RAD* predictions are accurate with a good compromise between recall and precision (85% and 93% respectively, see Table 1). Noteworthy, by selecting only highly scored variants (rank*>*0.5), almost all false positive predictions are discarded (precision of 99%) with a minor impact on the recall. Conversely, if one favors the recall, by relaxing all the constraints on the bubble model (branching mode -b 2), almost all simulated SNPs are recovered (except for the ones simulated at the first and last 3 base pairs of the loci, as explained in section 2.1), the highest reachable recall being here of 96%. By comparison with other tools, *DiscoSnp-RAD* obtains similar results as *STACKS* but with twice less missing genotypes, and can achieve the level of precision of *IPyRAD* but with a much better recall (note that we used *IPyRAD* with default parameters, which may include some filters that tune the results towards highly precise ones). Importantly, the number of clusters obtained with *DiscoSnp-RAD* is very close to the number of simulated loci (10,000), suggesting that predicted variants are well clustered by loci.

**Table 1.** RAD-Seq simulated results. “Rec.” stands for “Recall”, “Prec.” stands for “Precision”.Results are shown for three parameter sets of *DiscoSnp-RAD*: 1/ default mode (note that variants with rank<0.2 are discarded by default for RAD-Seq), 2/ in high recall mode with option `-b 2` instead of `-b 1`, and 3/ in high precision with a more stringent filter on the rank (rank>0.5).

|  | #<br>loci | SNP predictions |  | Missing |
| --- | --- | --- | --- | --- |
|  |  | Prec. (%) | Rec. (%) | genotypes (%) |
| *DiscoSnp++* default | 10,040 | 93.3 | 84.8 | 9.4 |
| *DiscoSnp++* high recall | 9,881 | 91.4 | 92.6 | 12.7 |
| *DiscoSnp++* high precision | 9,844 | 99.3 | 82.2 | 9.4 |
| *STACKS* | 9,845 | 91.1 | 88.4 | 23.4 |
| *IPyRAD* | 7,098 | 99.6 | 65.0 | 0.02 |

### 3.2 Results on real data

In this section, we present an application of the *DiscoSnp-RAD* implementation on ddRAD sequences obtained from the anthomyiid flies from the *Chiastocheta* genus. In this genus, classical mitochondrial markers are not suitable for discriminating the morphologically described species [5]. Although RAD-sequencing dataset phylogenies supported the species assignment [19], the interspecific relationships between the taxa could not be resolved with high confidence due to high levels of incongruences in gene trees [7, 19]. The dataset is composed of 259 sequenced individuals from 7 species. Results obtained on *DiscoSnp-RAD* were compared to the prior work of Suchan and colleagues, based on *pyRAD* analysis [19]. In addition, we provide a performance benchmark of *STACKS*, *IPyRAD* and *DiscoSnp-RAD* ran on this dataset.

#### Recovering all Chiastocheta species

Variant calling was run on the read1 files of the 259 *Chiastocheta* samples with *DiscoSnp-RAD*. Before the filtering, 115,920 SNPs and 34,703 INDELs were identified. After filtering, 2,553 SNPs and 1,838 INDELs, located in 1,314 clusters, were retained and are usable for population genomic approaches. This number of clusters, obtained on the reads 1, is coherent with the 1,672 loci from Suchan *et al.* [19], obtained on reads 1 and 2, but with markedly lower amount of missing data (23.5% as compared with 84% for *pyRAD*). Then, following the requirements of the STRUCTURE algorithm, only one variant per cluster was retained, resulting in a dataset composed of 1,314 variants including 861 SNPs. STRUCTURE successfully assigned samples to the seven species, both using SNPs and INDELs and SNPs only, consistent with the morphological species assignment and previously published results [19] (Fig.2). The assignment values represent the probability with which STRUCTURE assigns a sample to a cluster, depending on the information carried by the variants. Theoretically and in an extreme case, if all variants of a sample are completely differentiated, different from the others and specific to a cluster or species, the assignment will be 1. Using SNPs only, the assignment values are high with an average of 0.990 (sd 0.026) across samples and a minimum assignment of 0.779. These values are comparable to the assignment values obtained by Suchan *et al.* [19] with an average of 0.977 (sd 0.042) and a minimum of 0.685. INDELs increase the number of markers and increases slightly the average sample assignment with an average of 0.991 (sd 0.024). The phylogeny realized with RAxML on the 2,553 SNPs obtained after filtering, is congruent with the one obtained by Suchan and colleagues [19] (Fig.2). The internal branches separating the seven species are well supported by high bootstrap values.

**Fig. 2.**
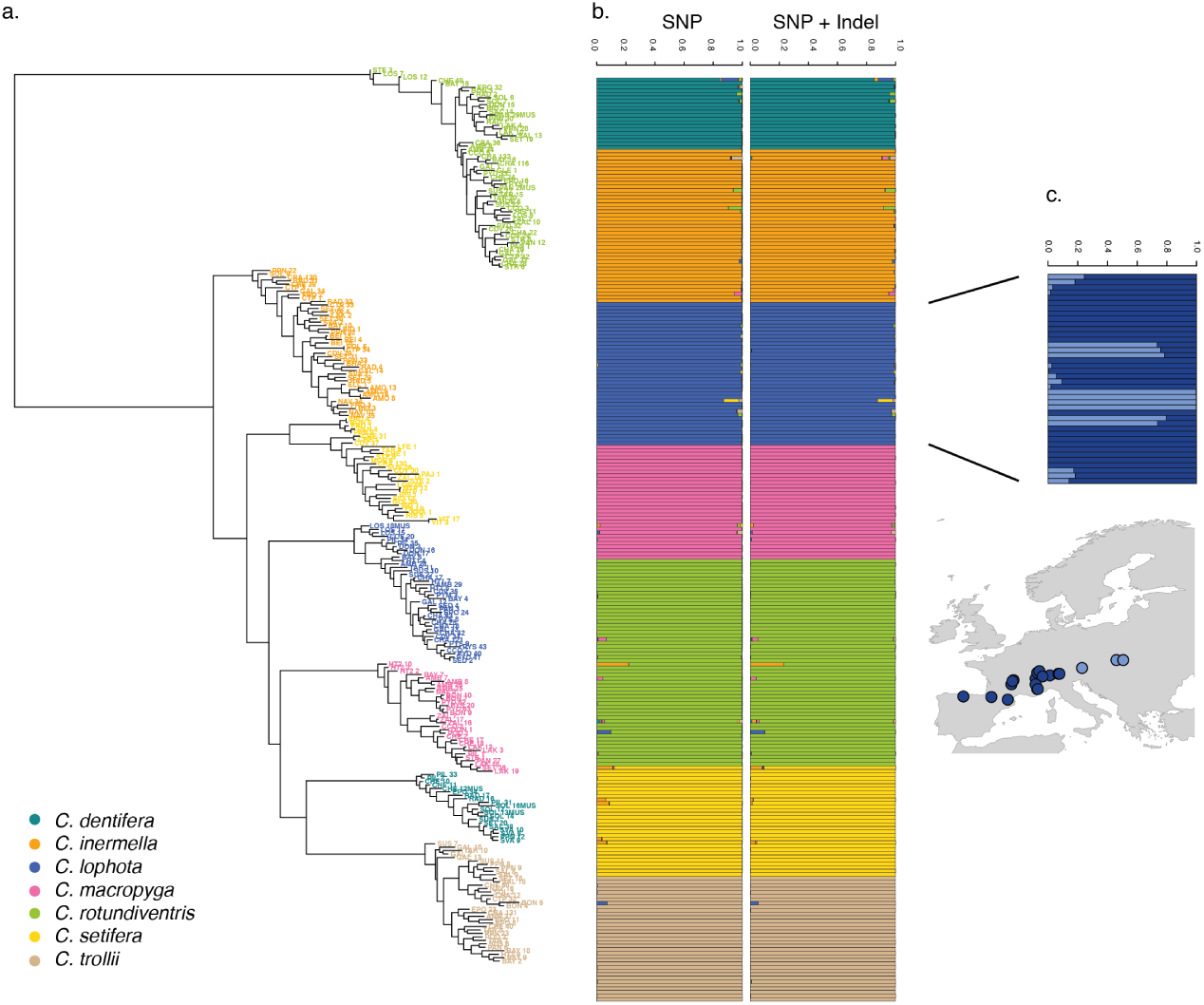
a. RAxML phylogeny realized on all variants predicted by *DiscoSnp-RAD*. STRUCTURE results obtained with SNP only and all variants on the seven *Chiastocheta* species. c. *C. lophota* sample structuration and their geographic distribution

#### Recovering phylogeographic patterns

To assess the utility of *DiscoSnp-RAD* dataset for investigating the intra-specific structuration, we focused the analysis on 40 samples from *C. lophota* species and adapted filters to extract informative variants at this evolutionary level. We obtained 822 SNPs and 2,141 INDELs by selecting one variant by locus extracted from 7,672 variants identified in this species. The STRUCTURE analysis of this dataset identified two populations and assigned 31 samples to one of them and 9 to the other (Fig.2). The assignment values are high with an average of 0.940 (sd 0.093). This structuration in two populations and the sample distribution within them is congruent with the geographic distribution which is the most frequent structuration factor observed in population genetics.

#### Breakthrough in running time

*DiscoSnp-RAD* run on the 259 *Chiastocheta* samples (30,063 Mbp overall) took about 4 hours. This comprises the whole process from building the dBG to obtaining the final filtered vcf file with for instance 1 SNP per locus. To compare the *DiscoSnp-RAD* performances with *STACKS* and *IPyRAD* on real data, we ran each of these tools on the 259 Chiastocheta samples (read 1 only) and measured running time and maximum memory usage. The difference is remarkable, *DiscoSnp-RAD* took more than 15 times less time than *STACKS* and *IPyRAD* to do the whole process (Table 2). Moreover, contrary to *DiscoSnp-RAD*, *STACKS* and *IPyRAD* should be run several times to explore the parameters which represent a considerable amount of time and memory. For instance, in Suchan *et al.* [19], *IPyRAD* was run with 5 different values of parameter, *DiscoSnp-RAD* being thus 89 times faster.

**Table 2.** RAD-Seq tool performance comparison on real data. ‡*STACKS* and *IPyRAD* run-times are shown for one parameter value, in practice several parameter values should be tested. For instance in [19], *IPyRAD* had to be run five times to determine parameters, the total time would be *≈* 22, 000*mn*, that is more than 15 days. All experiments were performed on a 20 cores cluster with 250GB of RAM.

|  | Time<br>(mn) | Memory<br>(MB) |
| --- | --- | --- |
| *STACKS* | 3,789<sup>‡</sup> | 15,264 |
| *IPyRAD* | 4,529<sup>‡</sup> | 18,950 |
| *DiscoSnp-RAD* | 252 | 8,114 |

## 4 Discussion

### DiscoSnp-RAD efficiency

*DiscoSnp-RAD* produced relevant results on ddRAD data from *Chiastocheta* species. Variants identified, SNPs and INDELs, allowed us to successfully i) distinguish the seven species based on the STRUCTURE algorithm, and ii) reconstruct the phylogenetic tree of the genus, congruent with the previously published one [19]. Moreover, on the intraspecific scale, we obtained geographically meaningful results within *C. lophota* species. The variants identified by *DiscoSnp-RAD* can be used to study the species or population structuration and could be used to investigate deeper the mechanisms at the origin of this structuration such as potential gene flow between populations or their demographic histories. Furthermore, the use of *DiscoSnp-RAD* presented considerable advantages in the run-time, and parameters choice, compared to other common *de novo* RAD analysis tools, as described below.

#### Run-time

The use of *DiscoSnp-RAD* dramatically decreased the overall time for discovering and selecting relevant variants, as compared to other tools. Moreover, *DiscoSnp-RAD* speed is less dependant on the number of reads and is not expected to increase quadratically with dataset size, as it is the case for pair-wise alignment based tools such as *STACKS* and *PyRAD*. This suggests that *DiscoSnp-RAD* will more easily scale to very large datasets, generated using high frequency cutting enzymes to obtain a dense genome screening, a deep sequencing to compensate sequencing variation or a large number of samples.

#### Easy parameter choice

Another substantial advantage of using *DiscoSnp-RAD* is the fact that parameters are not directly linked to the level of expected divergence of the compared samples. In fact, they impact the number and type of detected variants, but are not related to the subsequent clustering step. As a result, same parameters can be used whatever the type of analysis (for example, intra or inter-specific), contrasting with classical tools in which parameters govern loci recovering. Indeed, in *STACKS*, the parameters governing the merge of the stacks can compromise the detection of relevant variants if they are not adapted to the dataset used. Therefore, the authors recommend to perform an exploration of the parameter space before downstream analyses [13]. This is extremely time consuming, and may not always result in interpretable conclusions. In *IPyRAD*, the similarity parameter for clustering also impacts variant detection, and usually several values have to be tested to choose the best, as as exemplified by Suchan and colleagues who tested five different values [19].

#### By-locus assembly

*DiscoSnp-RAD* output is a vcf file including pseudo-loci information, that allows the application of standard variant filtering pipelines. One next objective is to recover loci consensus sequences, that could be used for phylogenetic analysis based on full locus sequences. This could be achieved by performing local assemblies per individual, from all bubbles contained in a cluster.

#### Potential applications

In many RAD-Seq studies using paired-end sequencing, the second read, i.e. the half of the sequencing effort, is only used to remove PCR duplicates and not exploited to detect variants. For example, current stable version of *STACKS* does not allow to exploit read 2 information in paired-end reads from experiments were fragments have been subjected to random shearing. Indeed, in such cases reads 2 do not start and finish at the same position, properly recovery of loci is therefore not possible. This problem does not exist when using *DiscoSnp-RAD*, and variants present in reads 2 can be called just like those present in read 1. In fact, whatever quality or INDELs or variations in coverage, all sequence information present in the reads is reflected in the dBG.

This ability of *DiscoSnp-RAD* to handle reads that do not necessary start at the same genomic position makes it particularly well suited to analyze the datasets produced by another group of genome-reduction techniques, namely sequence capture approaches [8]. In these techniques, DNA shotgun libraries are subject to enrichment using short commercially-synthesized [6] or in-house made [20] DNA or RNA fragments acting as ‘molecular baits’, that hybridize and allow separation of homologous fragments from genomic libraries. One of such promising approaches is HyRAD, a RAD approach combining the molecular probes generated using ddRAD technique and targeted capture sequencing, designed for studying old and/or poor quality DNA, likely to be too fragmented for RAD-sequencing [20]. In HyRAD, capturing randomly fragmented DNA results in reads not strictly aligned and covering larger genomic regions than RAD-Seq. Therefore RAD tools can not be used to reconstruct such loci, and the current analysis consists in building loci consensuses from reads, and then calling variants by mapping back the reads on it. The use of *DiscoSnp-RAD* should simplify this process in a single *de novo* calling step.

## Acknowledgments

This work was supported by the French ANR-14-CE02-0011 SPECREP grant. Authors thank Camille Marchet for her precious help on the clustering implementation. Computations have been made possible thanks to the resources of the Genouest infrastructures.

